# Molecular basis of ocean acidification sensitivity and adaptation in *Mytilus galloprovincialis*

**DOI:** 10.1101/2021.10.23.465493

**Authors:** Lydia Kapsenberg, Mark C. Bitter, Angelica Miglioli, Carles Pelejero, Jean-Pierre Gattuso, Rémi Dumollard

**Author notes:** shared first authorship (Lydia Kapsenberg, Mark C. Bitter).

## Abstract

One challenge in global change biology is to identify the mechanisms underpinning physiological sensitivities to environmental change and to predict their potential to adapt to future conditions. Using ocean acidification as the representative stressor, molecular pathways associated with abnormal larval development of a globally distributed marine mussel are identified. The targeted developmental stage was the trochophore stage, which is, for a few hours, pH sensitive and is the main driver of developmental success. RNA sequencing and *in situ* RNA hybridization were used to identify processes associated with abnormal development, and DNA sequencing was used to identify which processes evolve when larvae are exposed to low pH for the full duration of their larval stage. Trochophores exposed to low pH exhibited 43 differentially expressed genes. Thirteen genes, none of which have previously been identified in mussel trochophores, including three unknown genes, were expressed in the shell field. Gene annotation and *in situ* hybridization point to two core processes associated with the response to low pH: development of the trochophore shell field and the cellular stress response. Encompassing both of these processes, five genes demonstrated changes in allele frequency that are indicative of rapid adaptation. Thus, genes underpinning the most pH-sensitive developmental processes also exhibit scope to adapt via genetic variation currently maintained in the mussel population. These results provide evidence that protecting species’ existing genetic diversity is a critical management action to maximize the potential for rapid adaptation under a changing environment.

## Introduction

Anthropogenic environmental change is placing unprecedented stress on marine ecosystems. Species’ capacity to survive is determined by their ability to move to more favorable habitats, exploit phenotypic plasticity to persist in the new conditions, or evolve via selection for resistant genotypes (Hoffmann and Sgrò, 2011). While phenotypic responses to global change stressors are relatively straightforward to measure, understanding what drives such responses at a molecular level and predicting the potential for those mechanisms to evolve remain a significant challenge (Franks and Hoffmann, 2012). Here, we provide a first attempt to do so for the early development of a globally distributed marine bivalve in response to changes in seawater pH associated with ocean acidification.

Ocean acidification is a consequence of absorption of anthropogenic carbon dioxide (CO_2_) emissions by seawater and is particularly harmful to calcifying marine invertebrates, such as bivalves (Kroeker et al., 2013). Conditions of low pH, including low saturation of carbonate ions, challenges the formation of calcium carbonate shells and skeletons. Furthermore, CO_2_ dissolved in seawater can cross biological membranes and cause intracellular acidification (Venn et al., 2013), affecting enzyme and protein function (Casey et al., 2010). Despite a diversity of ocean acidification impacts on marine metazoans and a historic focus on calcifying species (Kroeker et al., 2013), a recent meta-analysis of transcriptomic studies revealed several common molecular responses that include, but extend far beyond, biogenic calcification (Strader et al., 2020). Still, mechanistic links between molecular and phenotypic responses, which ultimately drive population demographics (e.g., individual survival, development, and growth), remain largely unknown.

For many marine invertebrates, and particularly for molluscs (Gazeau et al., 2013), the larval stages are the most sensitive to changes in seawater carbonate chemistry (Gibson et al., 2011; Kroeker et al., 2013). Ocean acidification induces compounding impacts on the early development of marine mussels (Fig. 1). Briefly, exposure to acidified seawater leads to delayed development and both smaller and abnormal-shaped shells in D-veliger larvae (Kapsenberg et al., 2018; Kurihara, 2008; Waldbusser et al., 2015). Reduced shell size can be attributed to reduced carbonate saturation state in the calcifying space of the leading shell edge (Melzner et al., 2017). The consequence is a strong, and largely passive, dependence of calcification on external seawater chemistry (Kapsenberg et al., 2018; Melzner et al., 2017; Waldbusser et al., 2015). In contrast, the abnormal shape of D-veligers is driven by abnormal development of the underlying soft tissue caused by low pH conditions during the preceding trochophore stage (Kapsenberg et al., 2018). The primary developmental process at the trochophore stage is the development of the shell field (Fig. 1). The shell field is part of the larval epidermis and its cells secrete an extracellular, chitin-based, organic matrix within which calcium carbonate is precipitated to form the larval shell. The shell field develops from an invagination of ectoderm cells, followed by a pore-closure and secretion of the periostracum by a rosette of cells remaining at the surface (Kniprath, 1981, 1980). The deeper cells in the pore then evaginate back to the surface where, through mitotic divisions, the shell field expands and secretes the organic matrix of the larval shell under the larval periostracum. Interruption or abnormal progression of the shell field evagination is likely what gives rise to the abnormal D-veliger shell shape (Kapsenberg et al., 2018).

**Figure 1.**
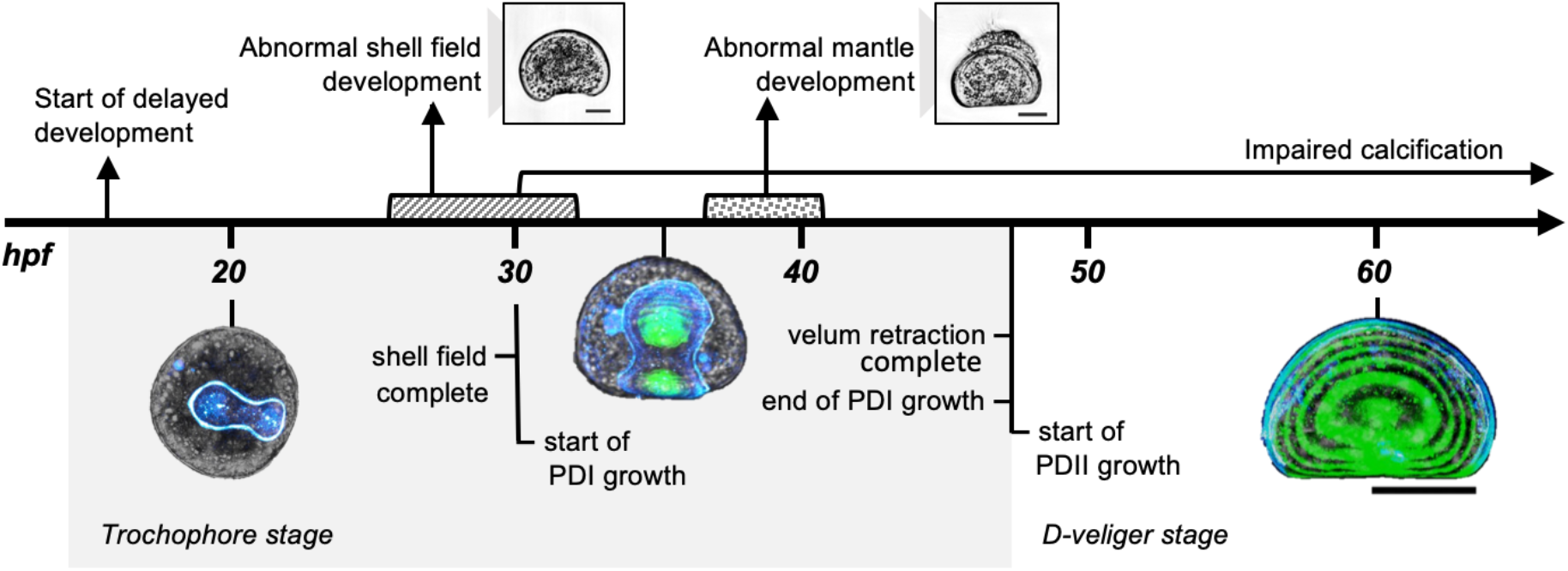
Ocean acidification impacts on early larval development. Chronologically, ocean acidification slows development, induces abnormal shell field development (hashed period, around 30 hpf) and mantle development (dotted period, around 40 hpf) during the trochophore stage which impacts D-veliger phenotype, and reduces carbonate saturation state in the calcifying space which impairs shell growth (30 hpf onward). Chronology represents development at 14°C; hpf is hours post-fertilization; PDI is Prodissoconch I; PDII is Prodissoconch II; colored larval images show the organic matrix secreted by the shell field (blue) and the calcified shell (green; scale bar is 50 μm), scale bar on D-veliger phenotypes is 30 μm. Figure is modified from Kapsenberg et al., 2018.

Fortunately, marine mussels are highly fecund which helps maintain substantial genetic variation in natural populations (Romiguier et al., 2014). Such variation may explain the diverse sensitivity to ocean acidification among different larval families (Frieder et al., 2017; Kapsenberg et al., 2018) and could contribute to rapid adaptation (Bitter et al., 2019). For example, previous research revealed that certain larval families are unaffected and exhibit entirely normal development in low pH conditions (Kapsenberg et al., 2018). Also, low pH conditions have shown to cause substantial changes to genetic variation in a genetically diverse larval population throughout development (Bitter et al., 2019).

To better understand the molecular underpinnings of pH sensitivity in larval development and the potential for this species to evolve and overcome ocean acidification stress, the aims of this study were to: (i) identify genes associated with the abnormal development of the shell field in trochophores of *Mytilus galloprovincialis* exposed to low pH conditions, and (ii) explore whether those genes that are physiologically responsive to low pH conditions harbor genetic variation that could drive rapid adaptation to ocean acidification. First, we performed genome-wide, RNA-sequencing to conduct differential gene expression analysis (‘RNA screen’ from hereon) in larvae exposed to control and low pH conditions during the trochophore stage (see ‘Exp. 3’ in Kapsenberg et al., 2018 for pH exposures and larval phenotypes). Differentially expressed genes (DEGs) were verified with qPCR, and the localization of larval expression of DEGs was identified using *in situ* hybridization (ISH) to provide potential links between the molecular and phenotypic responses to ocean acidification exposure. We then compared our list of DEGs to previous studies assessing bivalve gene expression patterns in response to low pH, as well as an independent list of genes shown to be subject to selection for low pH tolerance and reported by Bitter et al., 2019). This latter comparison (‘DNA screen’ from hereon) aimed to directly explore the likelihood for pH-sensitive biological processes to evolve in response to ocean acidification. Our results uncover a new set of candidate genes that may underpin the pH sensitivity of larval shell field development. Further, we provide evidence that genetic variation at these loci could facilitate rapid adaptation of *M. galloprovincialis* to ocean acidification.

## Results

### RNA screen: gene expression and localization in trochophores

Larval cultures exposed to low pH conditions (pH_Total(T)_ 7.4) during the trochophore stage exhibited 34% abnormal development, compared to 3% in larval cultures exposed to pH_T_ 8.1 during the trochophore stage. RNA-sequencing of trochophores exposed to low pH conditions revealed 38 differentially expressed genes (DEGs) compared to trochophores exposed to pH_T_ 8.1 that, after verification of each DEG’s DNA sequence, correspond to 38 different protein sequences (Table S1). Up-regulation in low pH was detected in 37 of the DEGs (logFC ranged from 0.32 to 1.25), including 15 genes with annotations. Up-regulation was confirmed by qPCR on 11 of the DEGs, 9 of which showed statistical significance (Table 1, Fig. S1).

**Table 1.**
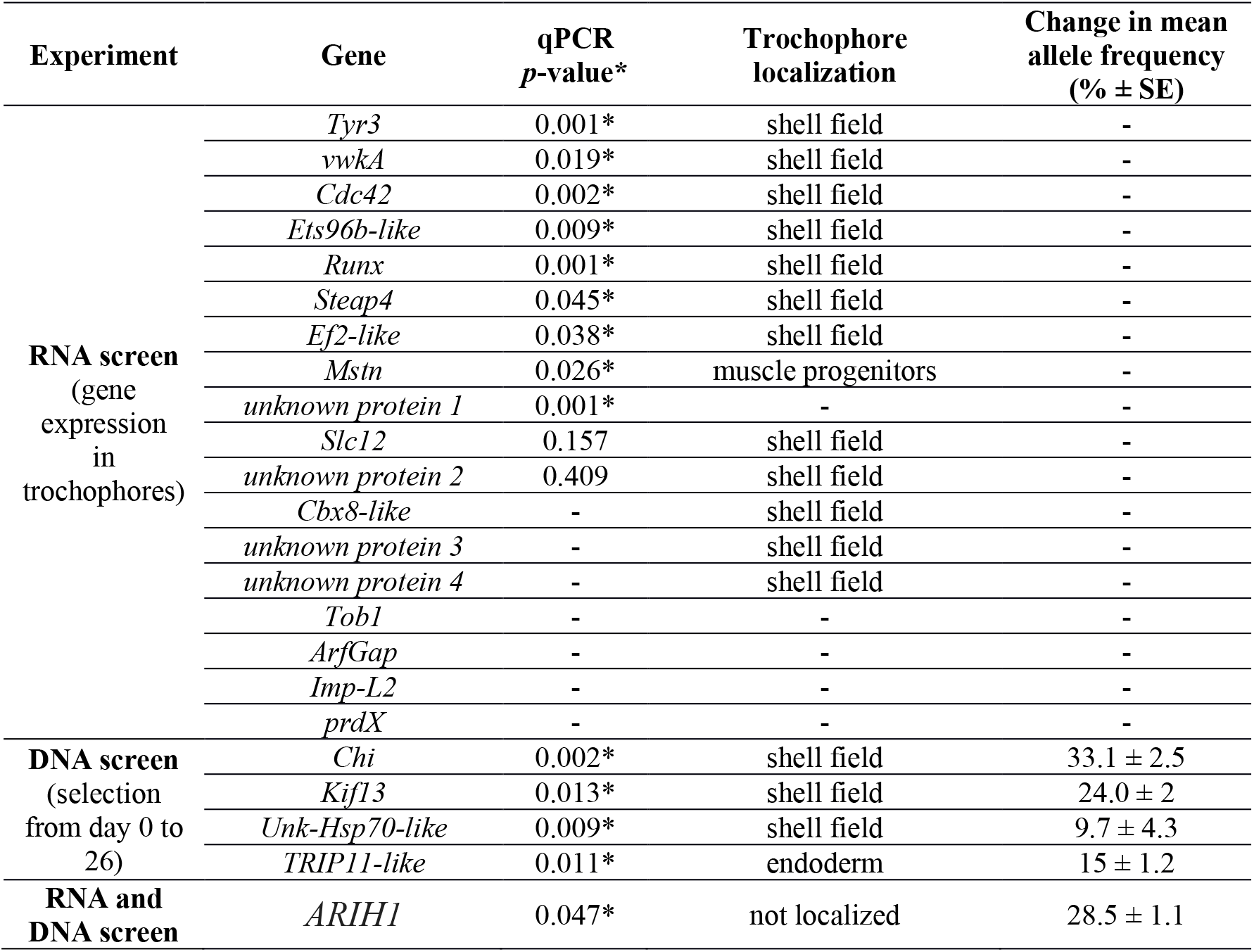
Annotated genes exhibiting up-regulation of expression during trochophore stage (‘RNA screen’) and allele frequency change from day 0 to 26 (‘DNA screen’) in larvae exposed to low pH conditions. *qPCR *p*-values on genes identified by the RNA and DNA screens both originate from qPCR performed on the same larval samples as used in the RNA screen.

*In situ* RNA hybridization on 14 of the DEGs demonstrated that 12 are expressed in the shell field: three genes, *Tyr3, vwkA, and Runx*, are known or suspected to be involved in larval shell morphogenesis (Middelbeek et al., 2010; Miglioli et al., 2019); seven genes (including *vwkA*) show localization in the shell field for the first time; and three genes encode unknown proteins (Table 1, Fig. 2). The remaining two DEGs that did not exhibit shell field expression were localized in the shell muscle progenitors (*Mstn*), or did not show conspicuous localization (*ARIH1*).

**Figure 2.**
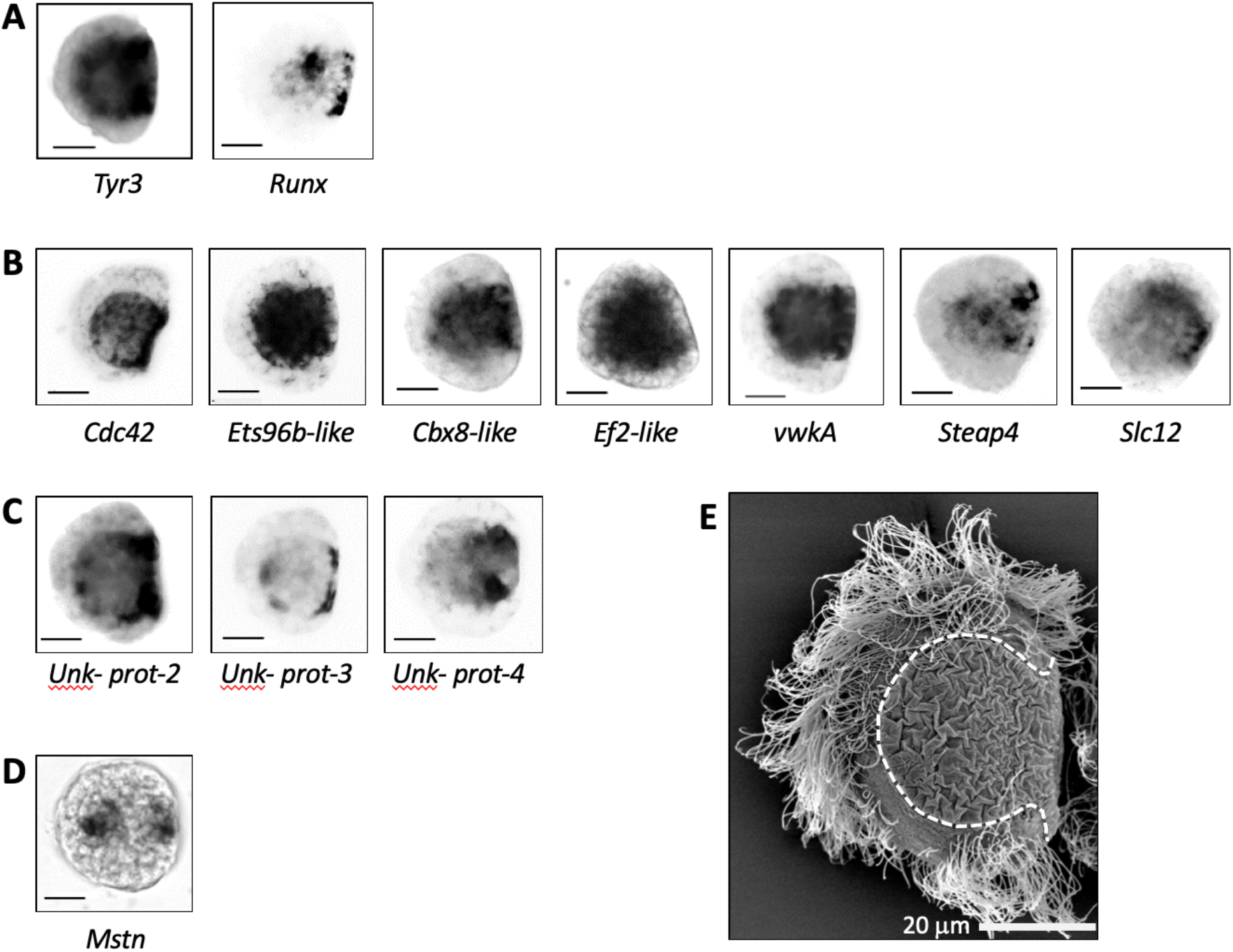
RNA *in situ* hybridization of genes in the mid-trochophore stage. Genes are those that exhibit differential expression, as determined by RNA-sequencing, when trochophores are exposed to pH_T_ 7.4. (A) Genes previously known to be expressed in the shell field. (B) Genes not previously known to be expressed in the shell field. (C) Unknown genes showing shell field expression. (D) Genes not showing shell field expression. Scale bar is 20 µm. (E) SEM image of a representative trochophore larvae; shell field is outlined (dashed). All imaged trochophore larvae were reared in pH_T_ 8.0 and fixed in the mid-trochophore stage (29 hours post-fertilization at 16.5 °C).

### DNA screen: Genes sensitive to pH that are prone to selection

To test whether molecular pathways sensitive to low pH conditions could lead to adaptation to ocean acidification, we compared the list of DEGs from the RNA screen to genes that exhibit signatures of selection during larval development reported by (Bitter et al., 2019). Briefly, *M. galloprovincialis* larvae were reared in pH_T_ 8.1 and pH_T_ 7.4 from the early embryo (4-cell stage) through settlement (day 43). Larvae were sampled on day 0, 6, 26, and 43 for reduced-representation genome sequencing using exome capture (hereafter referred to as DNA-seq). Changes in genetic variation within the larval population were quantified to identify genes undergoing persistent selection throughout the entire larval pelagic phase and settlement in low pH conditions (see Methods in Bitter et al., 2019 for more detail). This DNA screen resulted in 30 outlier loci mapping to annotated genes, only one of which, *ARIH1*, was differentially expressed in trochophore larvae exposed to low pH according to our initial RNA screen (Table 1, Fig 3A). However, due to the high statistical stringency (or batch effects) of this RNA screen, there are likely false negatives in that dataset (Pomaznoy et al., 2019). As such, we also explored the list of outliers from Bitter et al., 2019 to, *a priori*, identify genes that are also pH-responsive in other marine invertebrates. These genes were *Chi, TRIP11-like, Kif13, Hsp70*, and *Tyr1* (Beszteri et al., 2018; Bitter et al., 2021; Cummings et al., 2011; Glazier et al., 2020; Hernroth et al., 2011; Hüning et al., 2012; Kaniewska et al., 2015; Lardies et al., 2014; Li et al., 2017; Liu et al., 2020; O’Donnell et al., 2010; Padilla-Gamiño et al., 2013; Timmins-Schiffman et al., 2014; Wang et al., 2016). Note that for the transcript annotated as *Hsp70* in Bitter et al. (2019), we now refer to it as unknown-Hsp70-like (*unk-Hsp70-like*) given its unlikeness with the three known *Hsp70* in *M. galloprovincialis* genome upon close evaluation. Through qPCR, using the same samples as those used for the RNA screen, we found that *Chi, TRIP11-like, Kif13*, and *unk-Hsp70-like* were significantly up-regulated in response to low pH in trochophores, and so likely false negatives in the RNA screen (Fig. 3). ISH revealed that *Chi, Kif13* and *unk-Hsp70-like* are strongly expressed in the whole shell field (Table 1, Fig. 3), although only *Chi* has previously been known to be involved in larval shell development (Li et al., 2017; Liu et al., 2020). *TRIP11-like* is expressed in the endoderm. *Tyr1* was not detected by the RNA screen and showed no evidence of differential or localized expression at the trochophore stage (Fig. S2). Thus, *Tyr1* does not contribute to trochophore development, but likely instead supports larval survival in low pH conditions at a later developmental stage (Bitter et al., 2019).

**Figure 3.**
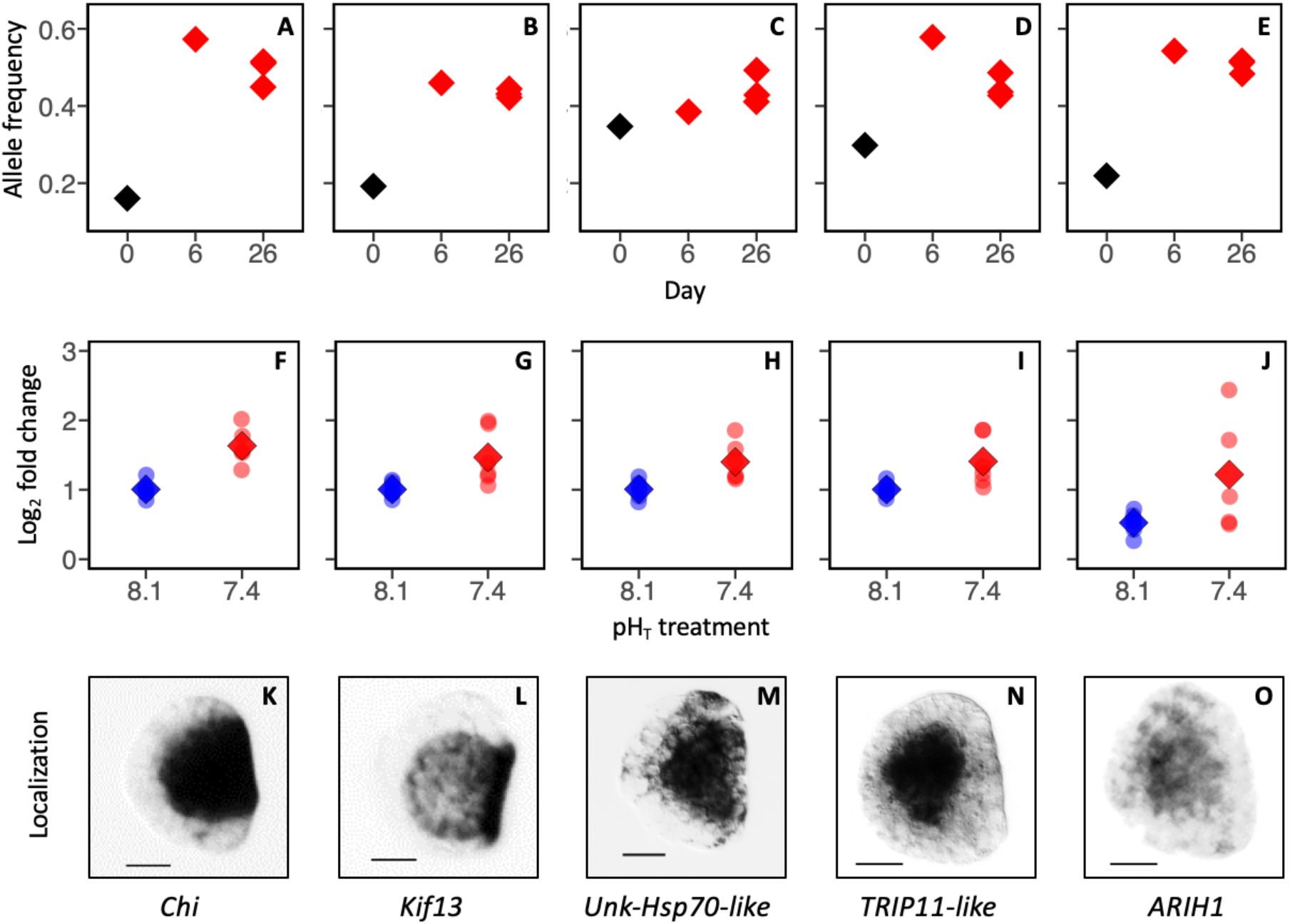
(A-E) DNA screen: frequency of selected allele at candidate genes on days 0 (*N* = 1) and 26 (*N* = 3) in larvae reared in pH_T_ 7.4. A constant was added to allele frequency values in panel (D) to present frequency shifts over the same range of values for all genes. (F-J) RNA screen using qPCR: log_2_-fold-expression of these genes in trochophore larvae exposed to pH_T_ 8.1 and pH_T_ 7.4 (faded circles indicate values individual replicate buckets, *N* = 6; diamonds correspond to treatment means). (K-O) Localization of these genes based on RNA *in situ* hybridization in trochophore larvae exposed to pH_T_ 8.0 (see Table 1 for localization).

To gain insight into the dynamics of low pH selection during early development, we examined changes in the frequency of genetic polymorphisms (alleles) during low pH selection from the embryo stage (day 0), through the trochophore to D-veliger transition (day 6), and during the pelagic feeding stage (day 26, Fig. 3). For all genes except *unk-Hsp70-like*, the most pronounced changes in allele frequency occurred between days 0 and 6 of development, which encompasses the trochophore stage when the course for abnormal development is set (Fig. 3). Specifically, the day 0 to day 6 change in allele frequency at these genes was 41% (*Chit*), 27% (*Kif13*), 28% (*Trip11-like*), and 32% (*ARIH1*). These large shifts in allele frequency were then maintained throughout the pelagic phase (Fig.3, Table 1).

## Discussion

Predicting the impact of global environmental change on species requires understanding both an organism’s sensitivity to changing conditions, and also how that sensitivity may be mitigated through time by natural selection on standing genetic variation and an associated increase in resilience (Hoffmann and Sgrò, 2011). Here, we targeted the precise developmental point during which low pH seawater induces abnormal development in *M. galloprovincialis*. By performing a differential RNA-seq analysis and reviewing published literature for target genes, a total of 43 pH-responsive genes were identified in trochophore larvae. *In situ* RNA hybridization performed on 18 of these genes revealed that 15 genes are expressed in the shell field. Of these 15 shell field genes, 13 genes were previously unknown to be expressed in the embryonic shell field of bivalves, which includes three genes encoding unknown proteins. Several pH-responsive genes show signatures of selection, indicating that these processes might support adaptation to ocean acidification. This includes three genes expressed in the shell field *Chi, Kif13* and *unk-Hsp70-like*, and one gene expressed in the endoderm, *TRIP11-like*, just under the shell field. While *Chi* is a major effector of shell morphogenesis, *unk-hsp70-like* and *ARIH1* may mediate cellular stress response.

### A core set of genes shed new light on shell field development

We previously suggested that the abnormal D-veliger phenotypes induced by low pH exposure are a result of altered development of the shell field during the preceding trochophore stage (Kapsenberg et al., 2018). In support of our hypothesis that abnormal larval development is driven by abnormal or incomplete restructuring of the ectoderm during formation of the shell field, we found a number of DEGs that are involved in cell proliferation and migration. These include *Cdc42*, transcription factors *Runx* and *Ets96b-like*, and proteins *Cbx8-like* and *E2f-like*, all of which exhibit expression in the shell field. *Cdc42*, a Rho subfamily GTPase is known to be a central regulator of cytoskeletal organization, which supports cell shape changes and cell migration during embryonic morphogenetic movements, similar to that involved in invagination/evagination of the shell field (Murali and Rajalingam, 2014). In early embryos, *Runx* function is required sometimes, if not always, for both the proliferation of progenitor cells as well as the differentiation of their descendants (Coffman, 2009, 2003), while *Ets96b-like* is an *Ets* family transcription factor known to interact with *Runx* during gene regulation (Arman et al., 2009). The oyster ortholog of Runx (also called *Runt*) is also known for regulating nacre formation in pearl oysters (Zheng et al., 2017) and is expressed in the shell field of their larvae (Song et al., 2019), but was not previously known to be pH responsive. *Cbx8* and *E2f* proteins are part of cellular machineries regulating cell cycle progression and have been linked to proliferation of cancerous cells (Kent and Leone, 2019; Xiao et al., 2019). Up-regulation of these five genes could point to the mechanisms by which ocean acidification causes the observed abnormal development of the ectoderm into the shell field (Kapsenberg et al., 2018).

Several DEGs that are expressed in the shell field appear to be involved in the development of the secreted, chitin-based organic matrix. For example, *Tyr3, Chi*, and *vwA* (von Willebrandt factor A), have previously been implicated in embryonic shell morphogenesis in marine bivalves: *Tyr3* is a major gene involved in chitin remodeling during the formation of the organic matrix, *Chi* is a well-known biomineralization gene expressed in trochophores and involved in chitin synthesis (Li et al., 2017; Liu et al., 2020). *VwkA* is a kinase phosphorylating vwA containing protein, which are major molluscan shell matrix proteins (Zhao et al., 2018). These results are also consistent with a study of the oyster *Crassostrea gigas*, which found that the expression of *Chi* and *Tyr* is altered when larvae are reared under low pH conditions (Liu et al., 2020). All three genes are strongly expressed in the entirety of the shell field (Fig. 2 and 3). *TRIP11-like*, a thyroid receptor interacting protein that encodes the Golgi microtubule-associated protein 210 (GMAP-210), is essential for trafficking of secreted proteins and skeletal development, and mutations in this gene have been shown to cause the rare skeletal disorder achondrogenesis type 1A (Roboti et al., 2015). Our study suggests, for the first time, a role for *TRIP11-like* in shell field formation. In conclusion, differential expression of these genes suggests that the formation of the larval organic matrix is highly pH-sensitive and contribute to the delayed development associated with low pH exposure.

In addition, two proteins involved in ionic regulation were up-regulated by low pH. *Steap-4* (six transmembrane epithelial antigen of the prostate 4) is a metalloreductase involved in iron and copper homeostasis (Ohgami et al., 2006). *Steap-4* could play a role in the formation of the organic matrix, as iron is one of the most important minor elements in the shells of bivalves (49-51), while copper ions are necessary for tyrosinase function (Nagai et al., 2007). *slc12* is a Na^+^-K^+^-2Cl^-^ cotransporter regulating chloride intake (Wang et al., 2009), which may be involved in the low pH responses via up-regulation of HCO_3_/Cl pumps during high CO_2_ exposure in bivalves (Zhao et al., 2018). Furthermore, three unknown DEGs exhibited shell field expression and provide new candidates for future gene function research.

Our screen also revealed Myostatin (*Mstn*), which is a growth factor that inhibits muscle growth in vertebrates, and has been shown to be up-regulated in response to environmental changes in adult clams (Morelos et al., 2015). To our knowledge, this is the first record of embryonic expression of *Mstn* in bivalves, and its up-regulation may also contribute to the observed developmental delay under low pH.

Finally, we found evidence for a cellular stress response in trochophores exposed to low pH, through the observed up-regulation of *unk-Hsp70-like* and *ARIH1*. Heat shock proteins (Hsp) serve as molecular chaperones that cope with the stress-induced denaturation of other proteins (Feder and Hofmann, 1999), and shifts either *Hsp70* expression or protein production have been observed in adult *M. galloprovincialis* (Bitter et al., 2021), red urchins (O’Donnell et al., 2009), oysters (Timmins-Schiffman et al., 2014; Wang et al., 2016), clams (Cummings et al., 2011), sea stars (Hernroth et al., 2011), and snails (Lardies et al., 2014). However, *unk-Hsp70-like* is not a *bona fide Hsp70* from *Mytilus*, but a completely novel gene that has never been characterized and further research is needed to ascertain its roles in embryonic development of bivalves. *ARIH1* is an Ariadne1-type E3 ubiquitin ligase similar to Parkin, which bind to the substrates of misfolded or damaged proteins (Flick and Kaiser, 2012) and suppresses cell death induced by unfolded protein stress (Imai et al., 2000). Ubiquitin ligases are involved in the low pH response in other marine invertebrates including oysters (Timmins-Schiffman et al., 2014), urchins (O’Donnell et al., 2009), corals (Kaniewska et al., 2015), and coccolithophores (Jones et al., 2013). In trochophores, *ARIH1* exhibits expression in the endoderm, indicating that ocean acidification induces stress in non-calcifying tissues. If, and how, cellular stress might contribute to abnormal tissue development in trochophores remains unknown.

### A core set of genes underlies adaptive potential to low pH

Accurately predicting the impacts of climate-related stressors on natural populations hinges upon identifying the extent to which adaptation may offset phenotypic consequences. Adaptation to environmental change can proceed rapidly when populations evolve via variation currently maintained within natural populations (Barrett and Schluter, 2008; Messer et al., 2016). The presence of adaptive genomic variation within *M. galloprovincialis* was first reported by Bitter et al., 2019, and those results have been leveraged here to assess allele frequency dynamics at specific genes with putative links to the phenotypic traits affected by low pH conditions. Based on Bitter et al., 2019, and other ocean acidification studies, we identified 5 genes, *Chi, Trip11-like, Kif13, unk-Hsp70-like*, and *ARIH1* that are pH-responsive and also exhibit substantial scope for rapid adaptation in *M. galloprovincialis*. These genes are involved in the development of the shell field (*Chi, Trip11-like, Kif13*) and the cellular stress response (*unk-Hsp70-like, ARIH1*).

Changes in allele frequencies at these genes from the embryo stage to the shell growth period (day 26) were substantial, ranging between 10 and 33%, indicating a strong selective advantage of those individuals harboring the adaptive allele in low pH conditions. Interestingly, for four of these genes (*Chi, Trip11-like, Kif13*, and *ARIH1*) the selected allele increased most dramatically between days 0 and 6, the period encompassing the trochophore stage. This verifies that developmental processes occurring within this time window, such as shell field development and successful transition to the D-veliger stage, are important for survival and adaptation (9, 18).

The only gene exhibiting linear changes in allele frequency throughout the sampling period was *unk-Hsp70-like*. While this gene was localized to the shell field, further research is needed to determine how it may participate in the bivalve stress response and to decipher its role in low pH adaptation. The cellular stress response is induced by low pH conditions in a range of marine taxa including other marine invertebrates with similar life-history characteristics to *M. galloprovincialis* (e.g. high levels of genetic diversity, relatively short generation times) (Strader et al., 2020). Thus, this pathway may similarly exhibit genetic variation that supports rapid adaptation in a range of putatively sensitive species.

It is important to note that the specific alleles analyzed here are not expected to be causal in driving differences in larval performance. Rather, they may serve as marker alleles closely linked to the genetic variants driving fitness differences between genotypes in low pH. Given that each of these genes exhibited differential expression in response to low pH during the trochophore stage, it is possible that causal variants occur in the cis-regulatory regions of these genes, and differences in survivorship are caused by genotype-specific patterns of expression at critical developmental points (Wittkopp and Kalay, 2012). Future tests of this hypothesis would more finely resolve the molecular underpinnings of low pH adaptation in marine bivalves.

## Conclusion

Given the focus of marine biology research on characterizing species sensitivity to climate-related changes, there is a need to expand research on understanding adaptation potential (Sunday et al., 2014). Through a series of independent screens and analyses, we identified a core set of genes associated with the pH-sensitivity of shell field development and found evidence for rapid adaptation thereof. Such adaptation could mitigate the anticipated negative impacts of ocean acidification. This study contributes new evidence to our increasing knowledge on how species could develop resilience to changing conditions (Bitter et al., 2019; Brennan Reid S. et al., 2019; Kapsenberg and Cyronak, 2019). Understanding how resilience could be bolstered will help inform management practices (Kapsenberg and Cyronak, 2019). For *M. galloprovincialis*, the results presented here suggest that protecting species’ existing genetic diversity is a critical management action in order to maximize the potential for rapid adaptation.

## Methods

### RNA screen (gene expression in trochophores)

*M. galloprovincialis* larvae were reared in a lab system under stable and variable pH treatments as described by Kapsenberg et al., 2018 (‘Exp. 3’ therein). Briefly, embryos from nine unique parental pairs were pooled and distributed across three replicate cultures (*N* = 3), over four treatments at 14.3 °C: (i) stable pH_T_ 8.1, (ii) variable pH_T_ 7.4, wherein embryos started development at pH_T_ 7.4 but pH was increased to pH_T_ 8.1 during the trochophore stage and reduced back to pH_T_ 7.4 during the early D-veliger stage, (iii) stable pH_T_ 7.4, and (iv) variable pH_T_ 8.1, wherein embryos started development at pH_T_ 8.1 but pH was decreased to pH_T_ 7.4 during the trochophore stage and increased back to pH_T_ 8.1 during the early D-veliger stage. Samples were grouped according to the pH exposure at the time of sampling, regardless of whether or not the pH treatment was fluctuating or static. This was done in accordance to phenotypic outcomes of those treatments and resulted in two groups (*N* = 6): (i) stable pH_T_ 8.1 and variable pH_T_ 7.4, which combined yielded 3% abnormal development, and (ii) stable pH_T_ 7.4 and variable pH_T_ 8.1, which combined yielded 34% abnormal development (see Fig. 3 in Kapsenberg et al., 2018).

Larvae were sampled at the mid-trochophore stage, 33 h post-fertilization (hpf). Approximately 60,000 larvae per culture vessel were preserved in TRIzol^®^ and frozen at -80 °C until RNA extractions, following manufacturer’s protocol. Quality of total RNA was verified on a NanoDrop and in an 1.5% agarose gel, and tRNA samples were aliquoted and stored at -80 °C. Samples were shipped on dry ice from France to the University of Chicago where tRNA quality was checked on a bioanalyzer (mean RIN score 9.5 ± 0.3, *N*=12) prior to library preparation (illumina® TruSeq Stranded mRNA). All samples were sequenced on a single lane of Illumina HiSeq4000 (100 bp, paired-end reads, average output of 6.4 megabases per sample) and a separate aliquot of tRNA was used for qPCR verification of candidate genes.

Raw reads were preprocessed using the publicly available expHTS pipeline to remove PhiX contamination, trim adapters and low quality read tails and polyA’s. Reads were mapped to a reference *M. galloprovincialis* transcriptome using SALMON (Moreira et al., 2015; Patro et al., 2017). The resulting counts for all samples were analyzed using custom R scripts and the edgeR library (Robinson et al., 2010). After filtering, a total of 48,000 transcripts were available for the identification of differentially expressed genes. Genes were identified as differentially expressed given an FDR corrected p-value < 0.05 and are hereafter termed ‘pH-responsive genes’. Annotation of differentially expressed genes was verified, and in some cases modified, from those provided in Moreira et al., 2014 by re-blasting sequences on NCBI. It was not possible to find potential orthologs of the several sequences that we decided to name “unknown protein”. These sequences are identified by their gene ID (Table S1) and can be found in NCBI protein database.

### qPCR verification

DEGs identified by transcriptomic analyses were validated using qPCR. qPCR was performed on tRNA samples from all 12 cultures (4 treatments, 3 replicates) produced in Kapsenberg et al., 2018. Quantity of tRNA samples previously extracted was determined via Nanodrop and quality was verified on a subset using a bioanalyzer (RIN score range 9.6-9.7, *N*=4). tRNA was converted to cDNA using the Applied Biosystems High-Capacity RNA-to-cDNA Kit, as per manufacturer instructions. qPCR reactions were performed on the Applied Biosystems Step One real-time qPCR machine, with a total reaction volume of 15 μL consisting of 7.5 μL SYBR Green PCR Master Mix (Applied Biosystems), 1 uL 10 μM primer (Table S2), 3 μL DNA, and 3 μL DI water per reaction. As with the transcriptome analysis, six replicate samples were run for each treatment. Controls lacking cDNA template were included in each reaction plate run to identify the presence of any non-target contamination. Amplifications were performed using the Applied Biosystems StepOnePlus real-time PCR system. Melting point assessment used to further determine the efficacy of each primer pair. Relative expression of target mRNAs was computed using the comparative CT method described in Schmittgen and Livak (2008) and is reported as the log_2_-transformed fold change in low pH (pH_T_ 7.4), relative to the control (pH_T_ 8.1) (Schmittgen and Livak, 2008). The significance of treatment on expression of each gene was determined using a permutational multivariate analysis of variance (PERMANOVA) in R with 999 permutations.

### In situ RNA hybridization

To identify the location of expression of pH-responsive genes within the larval body, primers for *in situ* hybridization (ISH) were designed from sequences when possible (Table S3) as described by Miglioli et al., 2019. Larval samples for ISH originate from an experiment independent of the transcriptomic and genomic approaches. Briefly, larvae from 4 unique male-female pairs were reared separately in static cultures (pH_T_ 8.0, 16.5 °C) and sampled at 29 hpf at which time larvae were in the mid-trochophore stage. Larvae were concentrated and fixed in 4% paraformaldehyde for 2 h at room temperature, rinsed with filtered seawater (FSW) three times, rinsed with 50% methanol for 15 min, and stored in 100% methanol at -20 °C. Samples were shipped on ice for preparation for *in situ* hybridization following methods described in Miglioli et al., 2019. Samples were only used if cultures of single male-female pairs yielded >90% normal development in pH_T_ 8.0.

### DNA screen (selection throughout larval development)

Data from Bitter et al., 2019 were used to identify whether or not DEGs in trochophores also exhibit adaptive variation for low pH tolerance. Briefly, the study crossed 16 males to each of 12 females and reared the resulting, genetically diverse, pooled larval population in pH_T_ 8.1 and pH_T_ 7.4 (17.2 °C) from the early embryo (4-cell stage) through settlement (day 43). Larval samples were collected for allele frequency estimation on days 0, 6, 26, and 43 and sequenced using exome capture, which targets the protein-coding region of the genome, facilitating the identification of genetic polymorphisms that are in or near functional regions of the genome. Outlier loci (i.e., genes under selection) were stringently identified as those genomic regions exhibiting significant shifts in allele frequency between the day 0 larval population and the larval population on all subsequent days of sampling, resulting in 30 outlier loci that mapped to functionally annotated genes (see Bitter et al., 2019 for details). These 30 genes were compared to the list of DEGs generated from the RNA screen, and an additional five genes from Bitter et al., 2019 that have previously been shown to be pH-responsive in other studies/species were tested for differential expression and location of expression in trochophores by qPCR and ISH, respectively.

Frequency of selected alleles were quantified in the starting embryo population (day 0), three days after the trochophore to D-veliger transition (day 6), and mid-way through the pelagic feeding stage (day 26) using pooled sequencing of the larval population (Fig. 4) (see Bitter et al., 2019 for further details).

## Acknowledgements

The authors thank Clàudia Aparicio for assistance with larval cultures and sample preparation for ISH and José Manuel Fortuño for assistance with SEM images used in Fig. 1, at the Institute of Marine Sciences (ICM) of the Spanish National Research Council (CSIC) facilities. We thank Soo Young Park and Graeme Bell of the University of Chicago for guidance and resources during qPCR assays. The unwavering assistance of Samir Alliouane and Lydia Besnardeau in the laboratory has greatly contributed to the success of the project.

## Author contributions

LK and MCB designed the study. LK, MCB, and AM performed the research and data analysis. LK, MCB, and RD wrote the manuscript with contributions from all authors. RD, JPG, and CP contributed input on experiments, resources, and data interpretation.

## Fundin

This research was funded by the US National Science Foundation (NSF; OCE-1521597 to LK) and European Commission Horizon 2020 Marie Skłodowska-Curie Action (No. 747637 to LK). MCB was supported by US Department of Education (Grant No. 200A150101) and NSF Graduate Research Fellowship (No. 1000198423).

## Supplementary Material

**Table S1.**
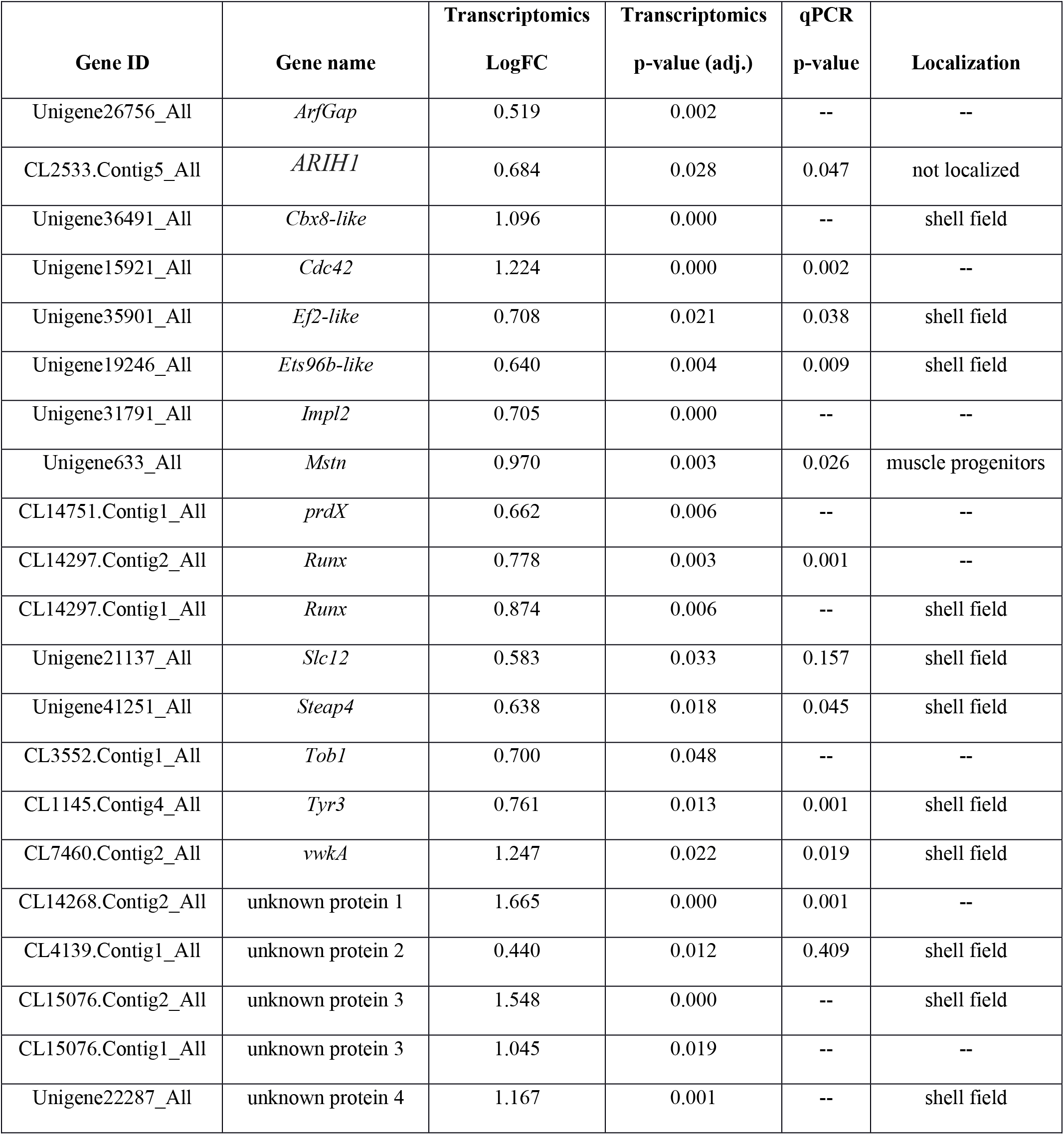
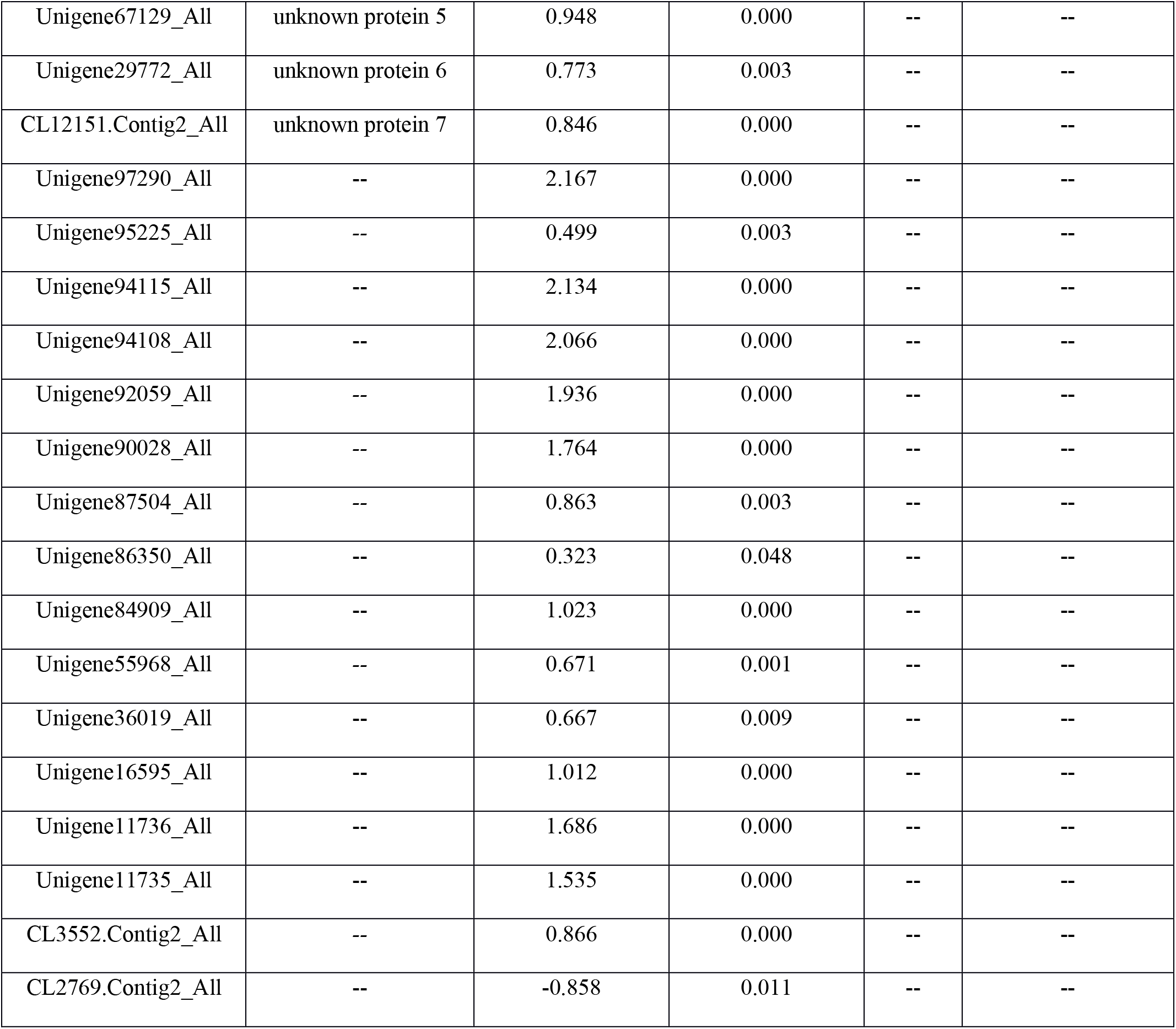
Differentially expressed genes in trochophore larvae exposed to low pH conditions at the time of sampling. Gene IDs correspond to the transcriptome published by Moreira *et al*. (1).

| Gene ID | Gene name | Transcriptomics<br>LogFC | Transcriptomics<br>p-value (adj.) | qPCR<br>p-value | Localization |
| --- | --- | --- | --- | --- | --- |
| Unigene26756_All | *ArfGap* | 0.519 | 0.002 | -- | -- |
| CL2533.Contig5_All | *ARIH1* | 0.684 | 0.028 | 0.047 | not localized |
| Unigene36491_All | *Cbx8-like* | 1.096 | 0.000 | -- | shell field |
| Unigene15921_All | *Cdc42* | 1.224 | 0.000 | 0.002 | -- |
| Unigene35901_All | *Ef2-like* | 0.708 | 0.021 | 0.038 | shell field |
| Unigene19246_All | *Ets96b-like* | 0.640 | 0.004 | 0.009 | shell field |
| Unigene31791_All | *Impl2* | 0.705 | 0.000 | -- | -- |
| Unigene633_All | *Mstn* | 0.970 | 0.003 | 0.026 | muscle progenitors |
| CL14751.Contig1_All | *prdX* | 0.662 | 0.006 | -- | -- |
| CL14297.Contig2_All | *Runx* | 0.778 | 0.003 | 0.001 | -- |
| CL14297.Contig1_All | *Runx* | 0.874 | 0.006 | -- | shell field |
| Unigene21137_All | *Slc12* | 0.583 | 0.033 | 0.157 | shell field |
| Unigene41251_All | *Steap4* | 0.638 | 0.018 | 0.045 | shell field |
| CL3552.Contig1_All | *Tob1* | 0.700 | 0.048 | -- | -- |
| CL1145.Contig4_All | *Tyr3* | 0.761 | 0.013 | 0.001 | shell field |
| CL7460.Contig2_All | *vwkA* | 1.247 | 0.022 | 0.019 | shell field |
| CL14268.Contig2_All | unknown protein 1 | 1.665 | 0.000 | 0.001 | -- |
| CL4139.Contig1_All | unknown protein 2 | 0.440 | 0.012 | 0.409 | shell field |
| CL15076.Contig2_All | unknown protein 3 | 1.548 | 0.000 | -- | shell field |
| CL15076.Contig1_All | unknown protein 3 | 1.045 | 0.019 | -- | -- |
| Unigene22287_All | unknown protein 4 | 1.167 | 0.001 | -- | shell field |
| Unigene67129_All | unknown protein 5 | 0.948 | 0.000 | -- | -- |
| Unigene29772_All | unknown protein 6 | 0.773 | 0.003 | -- | -- |
| CL12151.Contig2_All | unknown protein 7 | 0.846 | 0.000 | -- | -- |
| Unigene97290_All | -- | 2.167 | 0.000 | -- | -- |
| Unigene95225_All | -- | 0.499 | 0.003 | -- | -- |
| Unigene94115_All | -- | 2.134 | 0.000 | -- | -- |
| Unigene94108_All | -- | 2.066 | 0.000 | -- | -- |
| Unigene92059_All | -- | 1.936 | 0.000 | -- | -- |
| Unigene90028_All | -- | 1.764 | 0.000 | -- | -- |
| Unigene87504_All | -- | 0.863 | 0.003 | -- | -- |
| Unigene86350_All | -- | 0.323 | 0.048 | -- | -- |
| Unigene84909_All | -- | 1.023 | 0.000 | -- | -- |
| Unigene55968_All | -- | 0.671 | 0.001 | -- | -- |
| Unigene36019_All | -- | 0.667 | 0.009 | -- | -- |
| Unigene16595_All | -- | 1.012 | 0.000 | -- | -- |
| Unigene11736_All | -- | 1.686 | 0.000 | -- | -- |
| Unigene11735_All | -- | 1.535 | 0.000 | -- | -- |
| CL3552.Contig2_All | -- | 0.866 | 0.000 | -- | -- |
| CL2769.Contig2_All | -- | -0.858 | 0.011 | -- | -- |

**Table S2.**
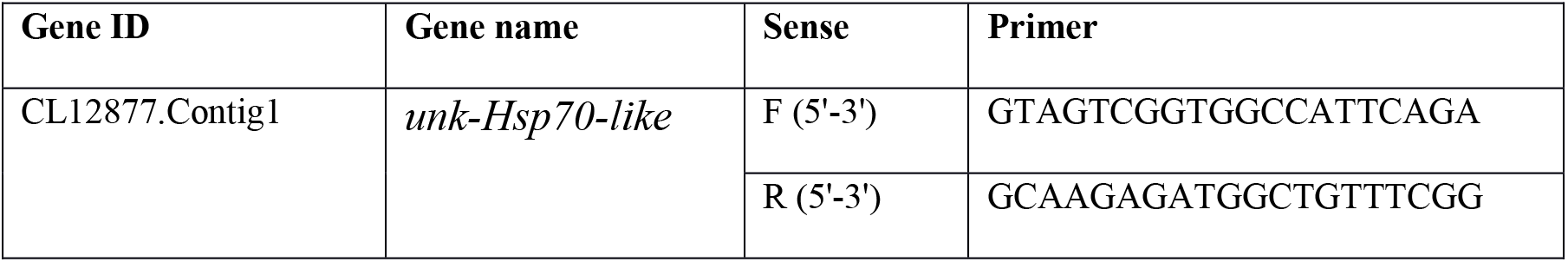
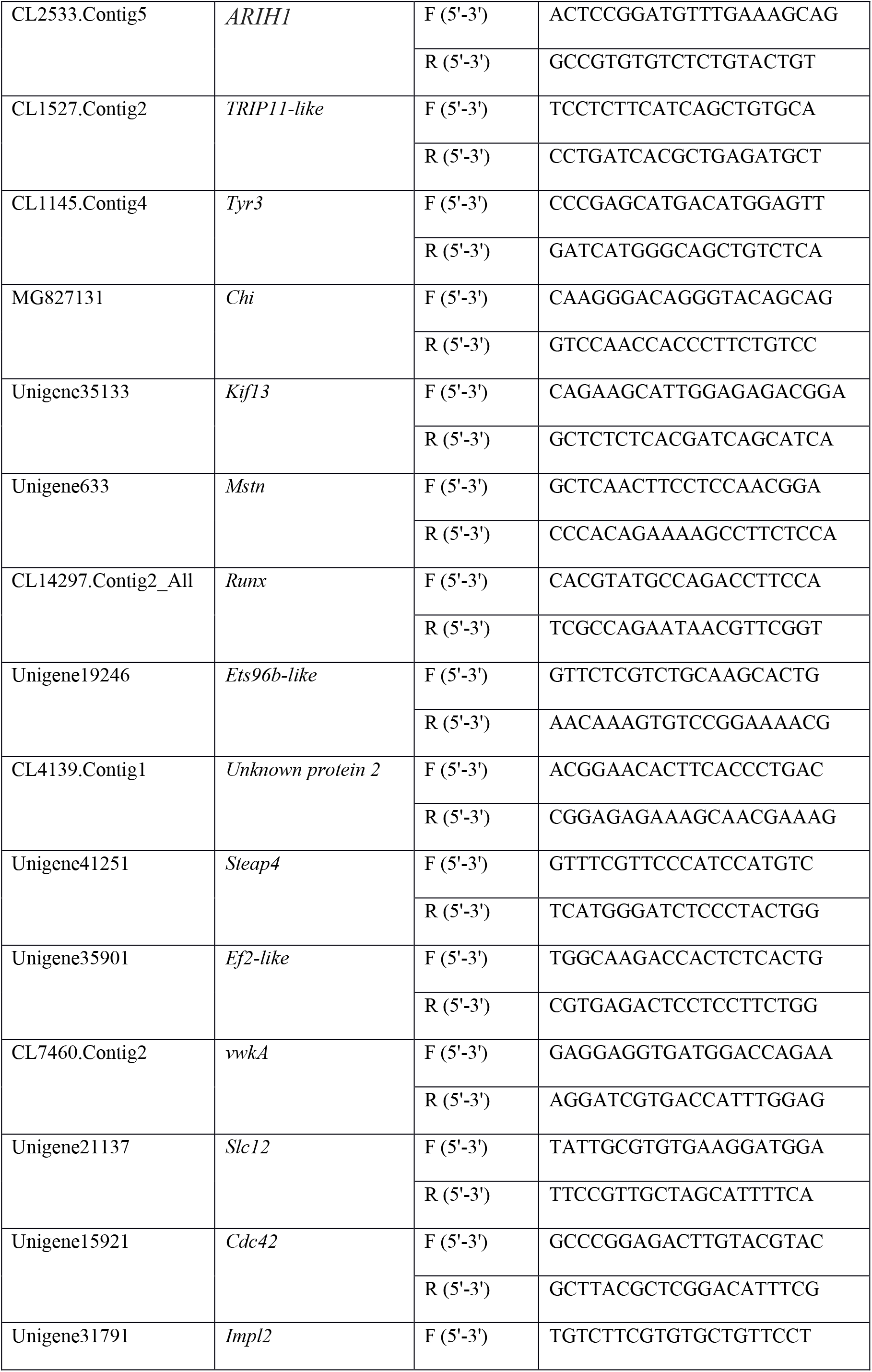
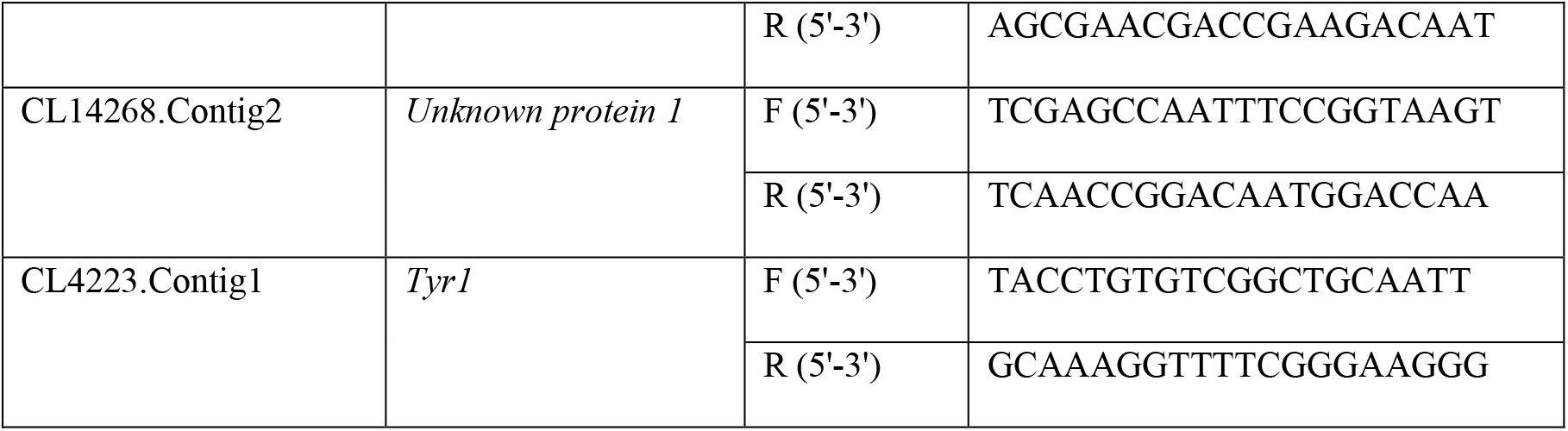
Primers used for qPCR. For the genes in which primers were designed directly from Gene ID corresponds to the associated gene ID in the Moreira *et al*. (2015) transcriptome. *Chi* required a longer sequence for qPCR primer design, so the *Chi* sequence from the Moreira *et al*. (2015) transcriptome was blasted to NCBI database to obtain sequences for primer design (the associated NCBI accession number is provided).

**Table S3.**
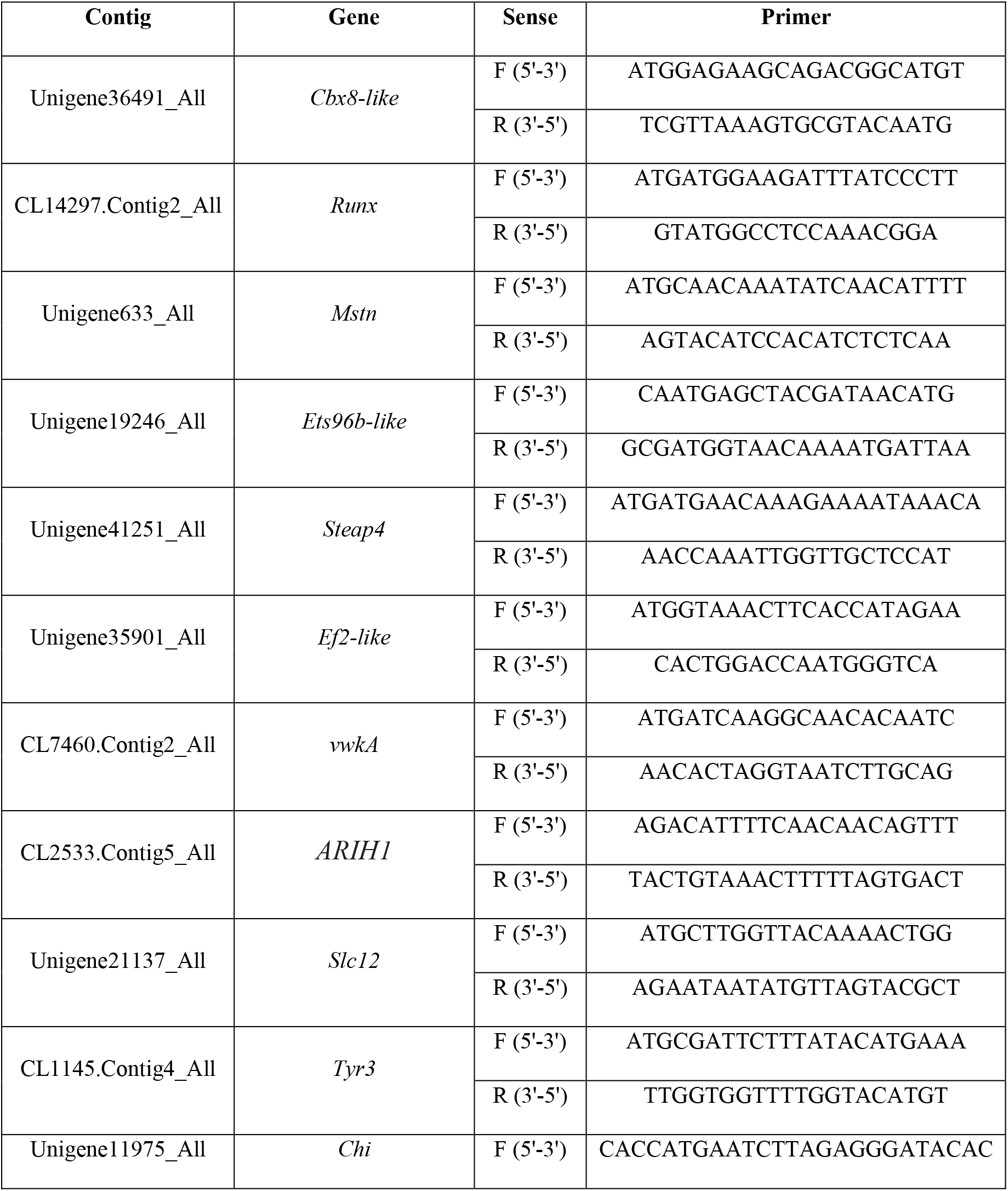
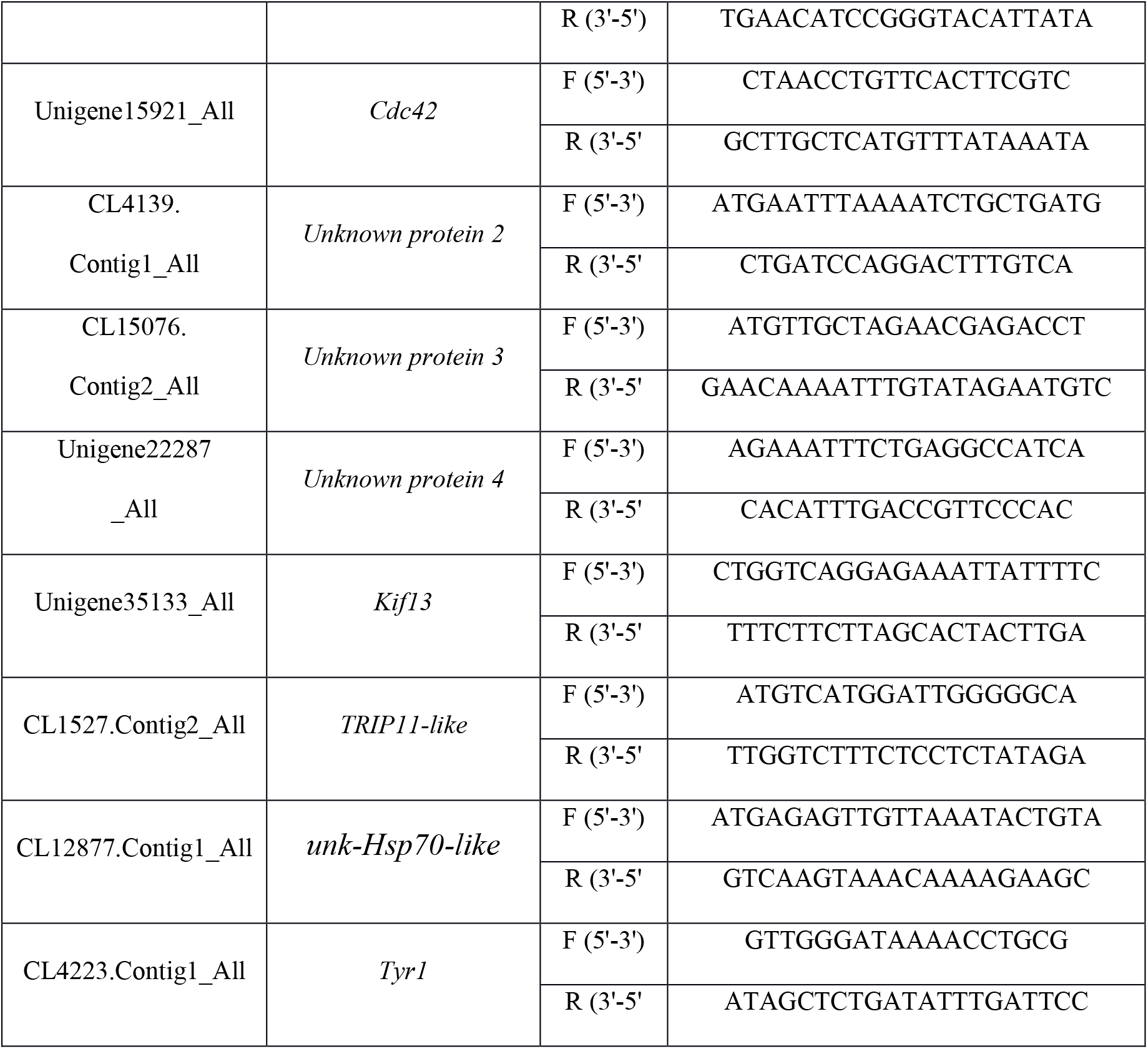
Primers used for in situ hybridization

---

**Figure S1.**
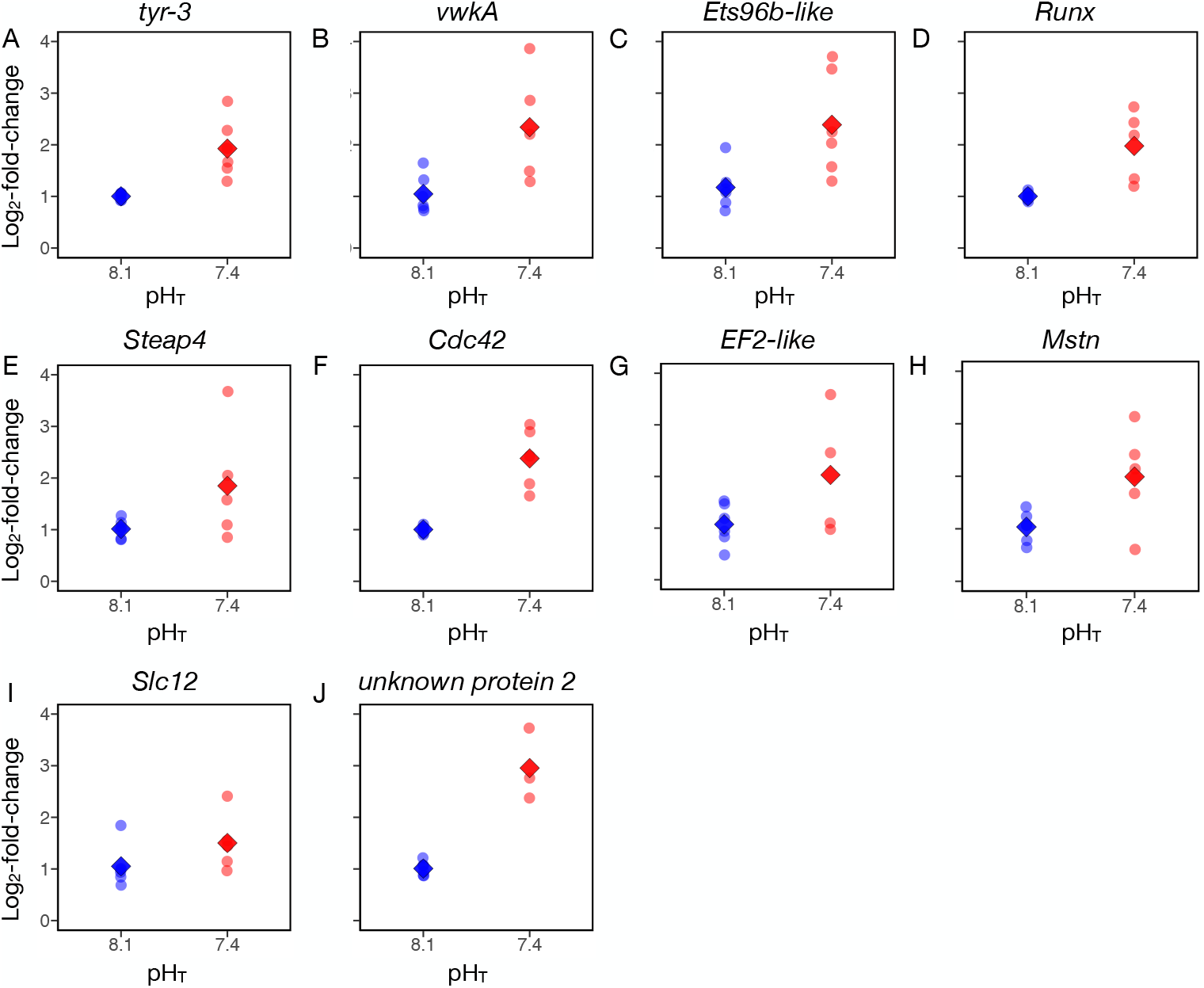
Log_2_-fold-expression at these genes identified as differentially expressed using RNA-sequencing, here determined via qPCR. Larvae were sampled at the trochophore stage at pH_T_ 8.1 (in blue) or pH_T_ 7.4 (in red, N = 6). Faded circles indicate values determined from individual replicate buckets, while diamonds correspond to treatment means.

**Figure S2.**
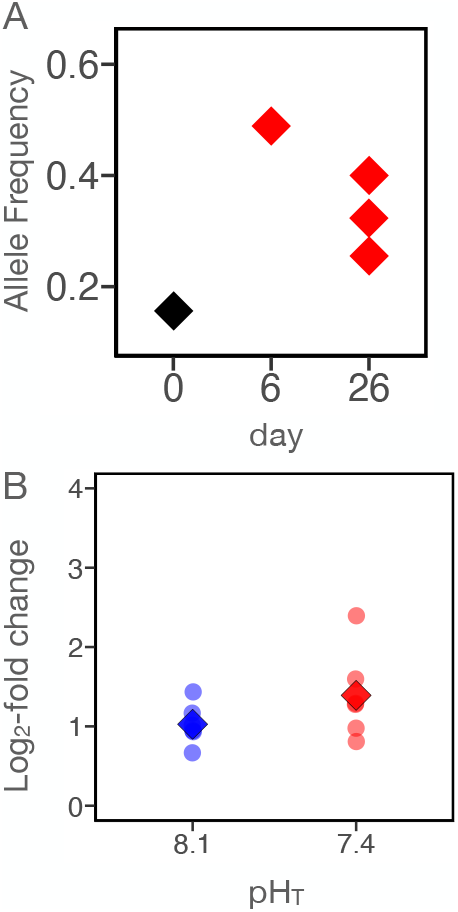
Patterns of expression in trochophore larvae, determined via qPCR (p > 0.05), and allele frequency change throughout larval development at *Tyr1*.

